# Cortical Representation of Pitch Perception in Mice

**DOI:** 10.1101/2025.09.04.674262

**Authors:** Jason W. Putnam, Abhay Kumar, Nasiru K. Gill, Jonathan Dinh, Franshesca Orellana Castellanos, Sofia Leusch, Sarah Vauhgn, Nikolas A. Francis

**Author notes:** These two authors contributed equally to this work (Jason W. Putnam, Abhay Kumar). **Corresponding Author Contact Information:** Nikolas A. Francis, 1210 Biology Psychology Building, College Park, MD, 20742-4415.

## Abstract

Pitch perception arises from temporal and spectral cues in sound. We hypothesized that mice rely on temporal cues because they have wide auditory filters, resembling humans with hearing loss or cochlear implants. Using computational modeling, behavioral assays, and widefield calcium imaging, we found that the structure of periodotopy in auditory cortex predicts how well mice recognize temporal pitch cues, establishing mice as a robust model for temporal pitch perception.

## Main Text

Pitch is the perception that sounds are organized from low to high on a scale derived from spectral and temporal auditory cues^1–11^. Pitch is critical for speech and music perception, yet individuals with hearing loss or cochlear implants struggle with pitch and rely more heavily on temporal cues^12–16^. Mice provide a powerful model for studying temporal pitch perception because they have high-frequency hearing and wide auditory filters^17,18^. Accordingly, periodic sounds with low fundamental frequencies (F0s) produce unresolved harmonics in the mouse auditory system and are represented as having a missing F0. Figure 1a illustrates how temporal pitch cues arise from unresolved harmonics. Importantly, mice can detect missing F0s in harmonic complex tones and perceive multi-harmonic communication sounds similarly to humans^19,20^. Moreover, mice are most sensitive to acoustic amplitude modulation rates less than several hundred hertz^21^, which is similar to the upper limit of F0 sensitivity for temporal pitch perception in humans^22^. We therefore asked if temporal pitch cues are salient to mice, under what conditions mice use temporal cues to discriminate F0s, and how cortical representations of F0 might reflect temporal pitch perception.

**Figure 1.**
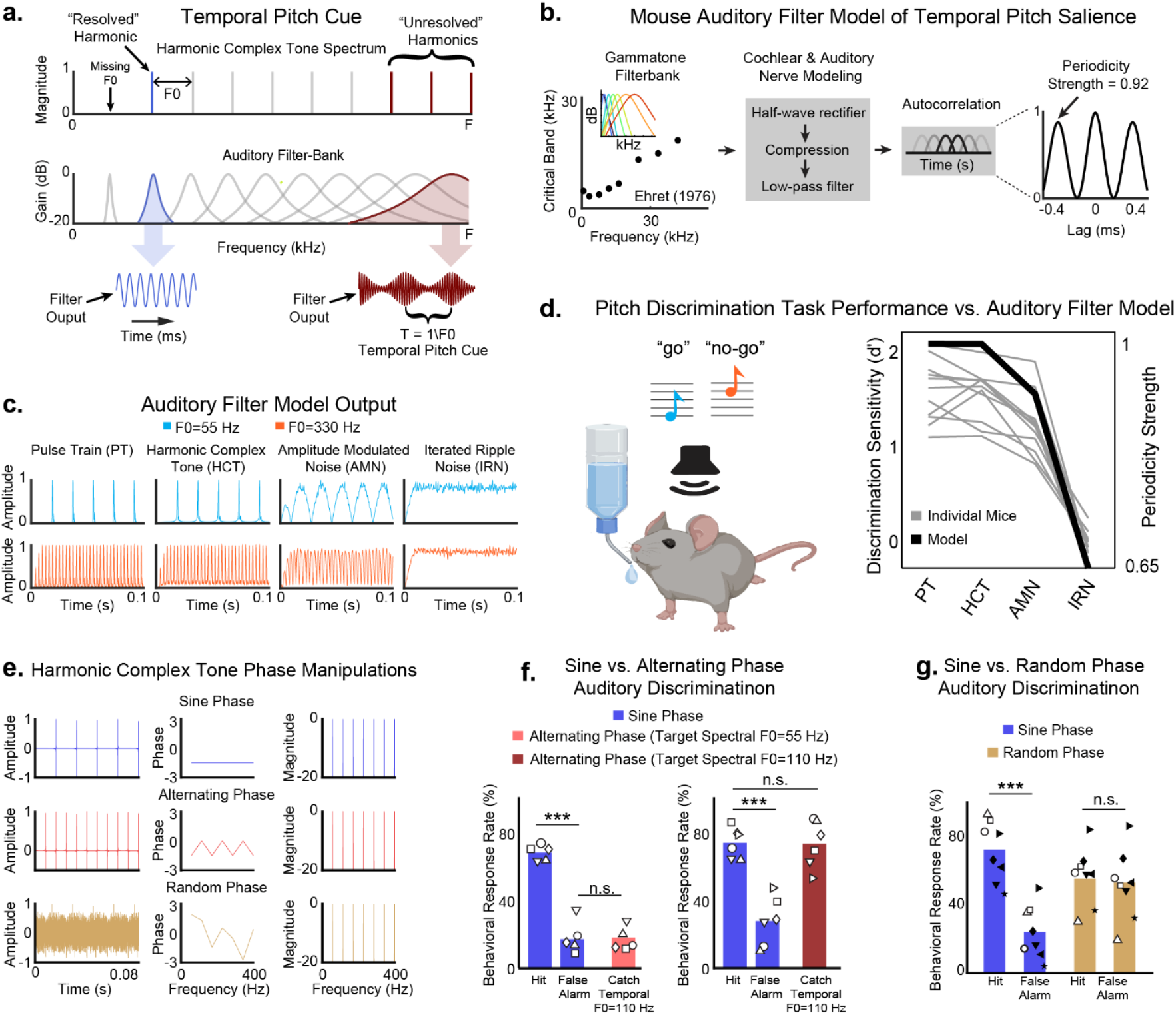
Temporal pitch perception in mice. **a. Illustration of how temporal pitch cues arise from unresolved harmonics.** When auditory filters have wide bandwidths, multiple harmonics are unresolved and fall within the same filter, causing periodic oscillations in filter output, giving rise to temporal pitch perception. [Adapted from Walker et al^11^]. **b. Mouse auditory filter model (see Methods)**. Autocorrelations were used to quantify periodicity strength in the output of a gammatone auditory filter model. **c. Auditory filter model output**. Model output for complex tones: pulse trains (PT), harmonic complex tones (HCT), amplitude modulated noise (AMN), and iterated ripple noise (IRN). **d. Behavioral testing of fundamental frequency (F0) discrimination, i.e**., **pitch discrimination**. *Left panel:* mice were trained on a go/no-go task to discriminate low vs. high F0s in PTs, HCTs, AMNs, and IRNs. *Right panel:* F0 discrimination sensitivity was highest for PTs and HCTs, intermediate for AMNs, and lowest for IRNs. Gray lines show individual mice. **e. Phase manipulations for harmonic complex tones. f. Sine-vs. alternating-phase F0 discrimination**. *Left panel:* mice tended not to respond during catch trials with alternating-phase harmonic complex tones with temporal F0=110 Hz matching non-target F0s. *Right panel:* mice tended to respond during catch trials with alternating-phase harmonic complex tones with temporal F0=110 Hz matching target F0s. **g. Sine vs. random-phase F0 discrimination**. Mice could not distinguish F0=55 vs. 330 Hz for random-phase harmonic complex tones. Bars show mean values. Symbols show individual mice. Filled and open symbols in figure 1g had spectral targets, F0=330 and 55 Hz, respectively. n.s. indicates that a difference is not statistically significant (p>0.05). Stars indicate statistical significance (bootstrap t-test, ***: p<0.001).

To estimate the salience of temporal pitch cues in mice, we developed a gammatone filter bank model^23^ of the mouse auditory periphery^24^, based on critical band measurements from Ehret^17^ (see Methods; figure 1b). Auditory filter output represents the pooled firing rate of auditory nerve fibers^6^. Our model consisted of a bank of bandpass filters, spaced at 10 Hz intervals from 4-48 kHz, followed by half-wave rectification and compression to simulate cochlear and auditory nerve processing^24–28^. For each input to the model, the autocorrelation of filter outputs was computed, and the height of the autocorrelation peak at the lag corresponding to 1/F0 was taken as the periodicity strength^2,3,27,29^ (PS; figure 1b). Periodicity strength indicates temporal pitch salience. We evaluated the model for four broadband (4-48 kHz) complex tones: pulse trains, harmonic complex tones, amplitude modulated noise, and iterated ripple noise (figure 1c). The model predicted that temporal pitch salience in mice would be highest for pulse trains (PS=1.00) and harmonic complex tones (PS=1.00), intermediate for amplitude modulated noise (PS=0.92), and lowest for iterated ripple noise (0.64) (thick black line in figure 1d).

We behaviorally tested the model predictions by training mice on go/no-go F0 discrimination tasks, i.e., pitch discrimination tasks (figure 1d-g). Mice initiated task trials by withholding licking for 5 s and were then presented with a 1 s complex tone. The complex tone for a given trial was broadband (4-45 kHz), presented at 75 dB SPL, and randomly selected from pulse trains, harmonic complex tones, amplitude modulated noise, or iterated ripple noise. Licking a waterspout within 2 s of a low F0 target (55 Hz) was rewarded as a “hit” (H), whereas licking after a high F0 non-target (330 Hz) was punished with a 30 s time-out as a “false alarm” (FA). F0 discrimination sensitivity was quantified using the signal detection theory metric, d’=z(H)-z(FA). d’≥1 indicates that mice were able to discriminate F0s. We found that F0 discrimination sensitivity was highest for pulse trains (d’=1.62±0.20) and harmonic complex tones (d’=1.59±0.16), intermediate for amplitude modulated noise (d’=1.25±0.19), while iterated ripple noise was significantly lower (d’=0.00±0.09, p<0.001, bootstrap t-test, Bonferroni correction). Task performance was strongly correlated (r=0.98±0.01) with model periodicity strengths, indicating that pitch salience predicts F0 discrimination sensitivity in mice.

Next, we directly tested whether mice rely on temporal or spectral cues to discriminate F0s. For a given F0, sine-, alternating-, and random-phase harmonic complex tones have identical spectral magnitude profiles but distinct temporal waveforms (figure 1e). Alternating-phase introduces waveform peaks at half the period of sine-phase tones, effectively doubling the perceived pitch arising from temporal cues. To determine when mice perceive temporal vs. spectral F0s, we trained a cohort of mice to discriminate sine-phase tones (spectral F0=55 Hz targets, 110 Hz non-targets; d’=1.48±0.30), interleaved with alternating-phase tones (spectral F0=55 Hz) during catch trials. Despite the 55 Hz spectral F0, mice withheld licking in response to alternating-phase tones, as if the perceived F0 was a 110 Hz non-target tone (figure 1f). Specifically, the false alarm rate during sine-phase (19.7±11.0%) was not significantly different than the response rate during alternating-phase catch trials (20.4±7.7%; p=0.81, bootstrap t-test).

We then trained a cohort of mice with the behavioral meanings of the sine-phase tones reversed (spectral F0=110 Hz targets, 55 Hz non-targets; d’=1.37±0.61) and also interleaved with alternating-phase tones (spectral F0=55 Hz) during catch trials. Here, mice treated alternating-phase tones as *targets*, again licking as if the perceived F0 were 110 Hz (figure 1f). Specifically, the hit rate during the sine-phase (74.4±8.9%) was not significantly different than the response rate during alternating-phase catch trials (73.8±11.8%; p=0.94, bootstrap t-test). When tested with random-phase perturbations that disrupt waveform periodicity, mice failed to distinguish F0s (d’=0.24±0.16) (figure 1g). Thus, we find that mice assigned behavioral meanings to harmonic complex tones based on temporal and not spectral pitch cues. Our results confirm that mice use temporal pitch cues to recognize low F0s.

Having established when mice listen to temporal pitch cues, we next examined the neural correlates of temporal pitch perception in auditory cortex using widefield calcium imaging in awake CBA × Thy1-GcaMP6s mice^18,30^ (figure 2). An increase in fluorescence during calcium imaging indicates an increase in the spike rates of local neuronal populations. Imaging experiments began with presentations of 75 dB SPL, randomly interleaved pure-tones, from 4-45 kHz. Subsequently, we presented pulse trains, harmonic complex tones, amplitude modulated noise, and iterated ripple noise, each at 75 dB SPL, broadband (4-45 kHz), and with F0s ranging from 20-1280 Hz.

**Figure 2.**
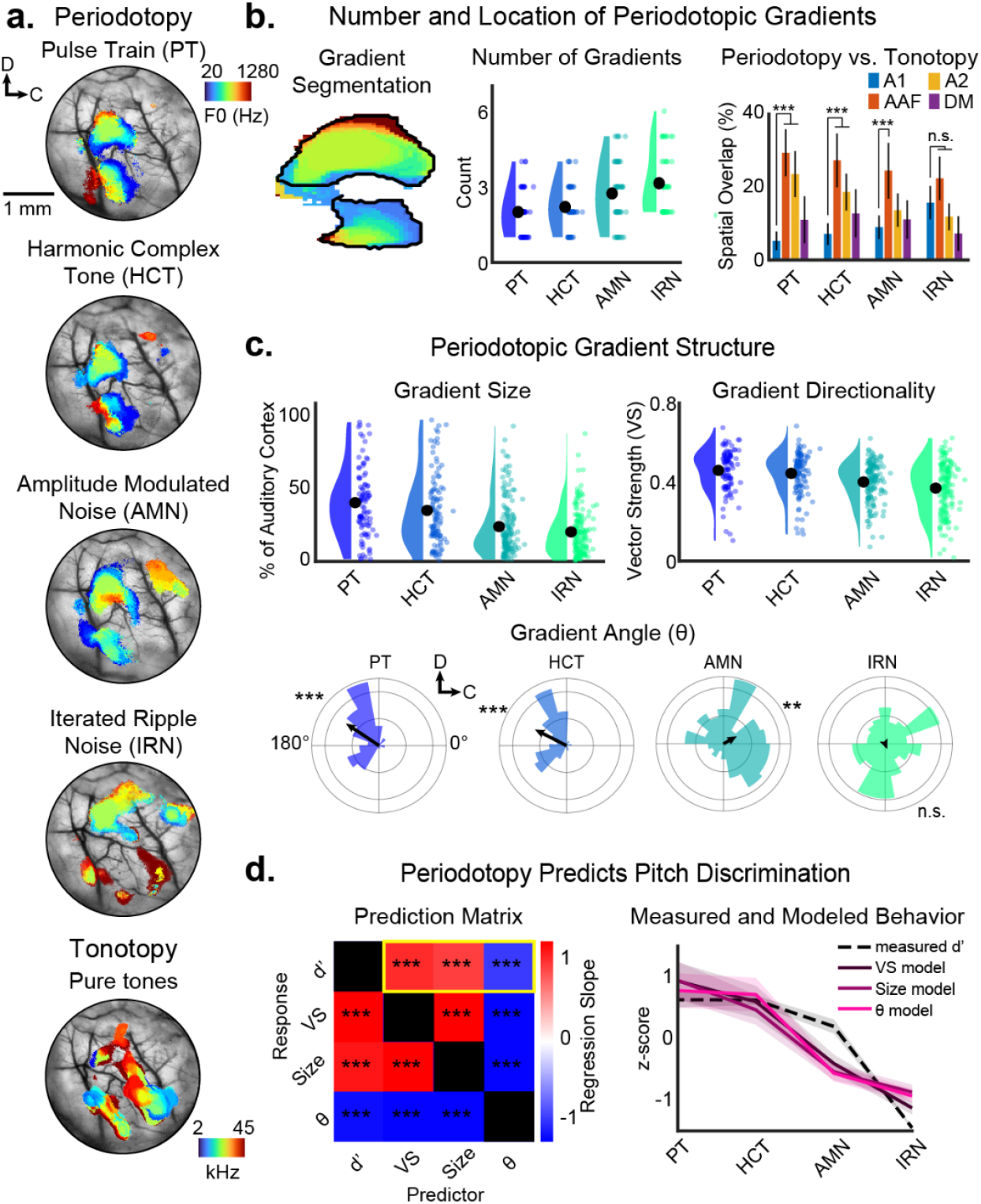
Periodotopy in auditory cortex (AC) predicts temporal pitch sensitivity in mice. **a. Example of periodotopy and tonotopy in a Thy1-GCaMP6s × CBA mouse**, mapped across AC using *in vivo* widefield imaging for 4 different complex tones: PTs, HCTs, AMNs, and IRNs. The color bar shows the range of tested F0s. **b. Number and location of periodotopic gradients**. We segmented periodotopic gradients in each imaging field of view (see Methods) from n=8 mice. The *middle* and *right panels* show the number of gradients found per stimulus type and the spatial overlap of periodotopy vs tonotopy, respectively. Each colored dot shows the count from an individual imaging session. We compare periodotopy against overlap with primary AC (A1), secondary AC (A2), the anterior auditory field (AAF), and the dorsomedial field (DM). Black dots show average values **c. Periodotopic gradient structure**. We quantified gradient structure using three metrics: vector strength (VS), size (% of AC), and angle. Each colored dot shows the value of an individual segmented gradient. **d. Periodotopy predicts pitch discrimination**. VS, gradient size (Size), gradient angle (θ), and behavioral sensitivity for F0 discrimination (d’) were used for pairwise comparisons using a repeated-measures linear regression analysis. *Left panel:* the matrix shows the estimated regression slope averaged across n=11 mice. The yellow box highlights the regressions for gradient structure as a predictor of d’. *Right panel:* z-scored comparison of modeled vs measured behavior. Shading shows 2 standard errors of the mean (SEMs). n.s. indicates that a difference is not statistically significant (p>0.05). Stars indicate statistical significance (bootstrap t-test, **: p<0.01, ***: p<0.001).

*In vivo* imaging revealed both periodotopy and tonotopy, i.e., topographic maps across auditory cortex of complex tone F0 and pure-tone frequency, respectively (figure 2a). Each complex tone produced multiple periodotopic gradients that were predominantly located outside of primary auditory cortex, in higher-order auditory fields including secondary auditory cortex (A2) and the anterior auditory field (AAF) (figure 2b).

To quantify the structure of periodotopy, we first segmented periodotopic maps into individual gradients (see Methods), then measured each gradient’s vector strength (VS; how much a gradient is oriented along a single direction), size (the percentage of auditory cortex covered by a gradient), and angle (θ; the direction a gradient is oriented in cortical space) (figure 2c). We found that the structure of periodotopy mirrored both behavioral task performance and model predictions. Vector strength and gradient size were largest for pulse trains (VS: 0.46±0.02; Size: 40.8±4.9) and harmonic complex tones (VS: 0.44±0.02; Size: 35.5±4.6%), intermediate for amplitude modulated noise (VS: 0.40±0.02; Size: 24.2±3.2) and smallest for iterated ripple noise (VS: 0.37±0.02; Size: 20.5±2.6). Gradient directionality followed the same trend, with pulse trains and harmonic complex tone gradients both pointing in a similar rostral-dorsal direction (θ=146.2±10.3° and 153.0±9.8°, respectively), amplitude modulated noise gradients pointing in a caudal-dorsal direction (θ=29.7±11.8°), and iterated ripple noise gradients being more diffuse, without a significant direction (θ=-68.6±12.4°, p=0.21, Omnibus test).

To quantify the relationship between periodotopy and F0 discrimination task performance, we regressed task performance (d’) against average gradient metrics (VS, Size, θ), separately for each behaviorally trained mouse shown in figure 1d (figure 2d). We also performed all pairwise regression comparisons for d’, VS, Size, and θ. VS showed the highest predictive power, with a slope of 0.84±0.10 and R^2^=0.81±0.07. Size had a slope of 0.77±0.10 and R^2^=0.67±0.08, whereas θ had a negative slope of −0.79±0.09 and R^2^=0.71±0.08 (p<0.001 for all predictors, with a Bonferroni correction for repeated-measures). Our results suggest that the salience of temporal pitch cues is encoded by the structure of periodotopic gradients in mouse auditory cortex.

Our study establishes mice as a robust model for temporal pitch perception. We show that mice rely on temporal pitch cues to discriminate F0s and that their discrimination sensitivity is predicted by the structure of periodotopic gradients in auditory cortex. Similar results have been observed across mammals. In marmosets, neurons in an anterior lateral region of auditory cortex are sensitive to the salience of temporal pitch cues^8^. In macaques, neurons in primary auditory cortex are tuned to temporal pitch cues^31^. In humans, neurons encoding temporal pitch cues have been found throughout Heschl’s gyrus, with stronger representations observed in higher-order auditory cortical fields^31–36^. This parallels our observation that periodotopy in mice is stronger in non-primary regions such as A2 and AAF. Indeed, periodotopy appears to occur widely across species, including cats, ferrets, gerbils, and humans, though the exact structure of periodotopy varies across species^37–41^. Thus, our findings align with a broader pattern in which temporal pitch cues are preferentially represented in higher-order auditory fields and expressed as maps across cortex. The existence of periodotopy suggests that place codes for F0 may provide a functional advantage in the cortical processing underlying pitch perception. Taken together, our results establish a comparative framework for investigating the neural basis of temporal pitch perception in mice and lay the groundwork for future studies of how cortical circuits transform sensory information into perceptual experience.

## Acknowledgements

This research was funded by NIH R21DC017829 (NAF), NIH T32DC000046 (JWP), and a UMD Brain and Behavior Institute seed grant (NAF). NAF designed the experiments, experimental software, and hardware. JWP, NKG, JD, FOC, SL, and SV performed experiments. NAF designed and implemented the auditory filter model. NAF and AK analyzed data and created figures. JWP, AK, NKG, JD, FOC, SL, SV, and NAF wrote the manuscript. The authors thank Dr. Catherine Carr for comments on initial drafts of this manuscript.

## Online Methods

### Auditory Filter Model

To simulate cochlear and auditory nerve processing in mice, we implemented a gammatone auditory filter bank model in MATLAB. Our approach captures auditory transformations relevant for temporal pitch processing in mice, but omits detailed mechanisms contained in models such as the Zilany et al (2009) model of the auditory periphery^24^. Four types of periodic sounds were synthesized for model analysis: pulse trains (PTs), harmonic complex tones (HCTs), amplitude modulated noises (AMNs), and iterated ripple noises (IRNs). All stimuli were generated using a 200 kHz sampling rate, tapered with 5 ms onset and offset ramps, normalized to unity magnitude, and band-limited between 4-48 kHz. Fundamental frequencies (F0s) were tested at 55 Hz and 330 Hz. HCTs were synthesized by summing sine-phase pure-tones, with harmonic frequencies spaced at integer multiples of the F0, beginning at F0. PTs were synthesized by concatenating impulses across silent temporal intervals of 1/F0. AMNs were synthesized by 1/F0 temporal envelope modulation of broadband noise at 100% depth. IRNs were synthesized using the delay-and-add method^29^ for a broadband noise token, using 5 iterations, and the delay set to 1/F0.

The model cochlea was represented by a bank of fourth-order zero-phase gammatone filters with center frequencies spaced every 10 Hz from 4-48 kHz. Filter bandwidths were set according to mouse behavioral critical band (CB) measurements reported by Ehret^17^ (CBs: 3.9, 4.4, 6.4, 7.9, 15.7, 17.7, and 21.9 kHz for characteristic frequencies of 5, 10, 15, 20, 30, 40, and 50 kHz, respectively), using a linear fit to interpolate values across frequencies. The impulse response of each gammatone filter was convolved with the input stimulus, yielding a band-limited waveform at each cochlear channel. The output of each gammatone filter was half-wave rectified to capture only positive-going responses, reflecting the asymmetric transduction of inner hair cells and because auditory nerve fibers fire only to positive deflections of the basilar membrane. A compressive non-linearity 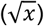 was then applied to the rectified signal to approximate the saturation and level compression introduced by cochlear mechanics and hair cell transduction. Finally, each channel was then low-pass filtered with a 6th-order Butterworth filter at 1 kHz to simulate the limits of phase locking in the mouse auditory nerve. The spectral profile of the auditory response was obtained by taking the maximum amplitude across time for each channel and normalizing by the channel’s peak response.

To quantify temporal pitch salience, we computed the autocorrelation function of the filter output within each channel. For each stimulus, the periodicity strength (PS) was defined as the autocorrelation function peak located at the lag corresponding to the fundamental period (1/F0). We then averaged across channels to obtain a single PS value per stimulus. The model is available here: https://bitbucket.org/FrancislabUMD/putnam_and_kumar_et_al_2025/src/main/

### Animals

All procedures were approved by the University of Maryland Institutional Animal Care and Use Committee. We used a total of N=29 mice (14 male, 15 female, aged 13-35 weeks) in our experiments. The mice were F1 offspring of CBA/CaJ mice (The Jackson Laboratory; stock #000654) crossed with transgenic C57BL/6J-Tg (Thy1GCaMP6s)GP4.3Dkim/J mice^30^ (The Jackson Laboratory; stock #024275). We used the F1 generation of crossed mice because they have both GCaMP6s expression in excitatory neurons and healthy hearing at least 1 year into adulthood^18^. 21 mice were used for behavioral experiments. 8 mice were used for imaging. 4 mice were behaviorally trained before imaging. All mice were housed under a reversed 12 h-light/12 h-dark light cycle.

### Acoustic Stimulus Presentation

During behavioral and imaging experiments, we presented awake mice with sounds from an ES1 free-field speaker (Tucker-Davis Technologies) connected to an ED1 amplifier (Tucker-Davis Technologies). The speaker was calibrated *in situ* to give a flat frequency response in the 1-80 kHz range. Our acoustic stimulus set included PTs, HCTs, AMNs, and IRNs. Stimuli were synthesized in MATLAB software (Mathworks) and output at 75 dB SPL from a National Instruments board (NI-6211, 200 kHz sampling rate) to the ED1 amplifier. Each stimulus had 5 ms and 495 ms raised-cosine attack and decay ramps, respectively. HCTs were synthesized by summing pure-tones spaced at integer multiples of the F0, beginning at F0. All components were typically in sine-phase, except during phase manipulation experiments, in which components were either (1) all in sine-phase, (2) alternating between sine- and cosine-phase for even and odd harmonics, respectively, or (3) set to a random phase uniformly distributed between −pi and pi. PTs were synthesized by concatenating impulses across silent temporal intervals of 1/F0. AMNs were synthesized by 1/F0 temporal envelope modulation of broadband noise at 100% depth. IRNs were synthesized using the delay-and-add method for a broadband noise token, using 5 iterations, and the delay set to 1/F0. All broadband sounds were bandpass filtered between 4-45 kHz.

### Behavioral Testing

F0 discrimination was tested using a custom training system based on a previous design^42^. Each operant chamber contained a TDT ES1 speaker calibrated *in situ* and a lick-sensitive waterspout. Water restricted mice were trained to lick a waterspout in response to hearing a target F0, and to avoid licking the waterspout after hearing a non-target F0. PT, HCT, AMN, and IRN presentations were randomly interleaved, and each type of complex tone was presented at target and non-target F0s. Each trial of the task was 4 s long with a randomized 5-9 s inter-trial interval. To initiate trials, the mouse refrained from licking for 5 s. Trials began with a 1 s silent period, followed by the 1 s stimulus, and ended with a 2 s silent period. If the mouse licked the waterspout between the sound onset and the end of the trial, i.e., a Hit (H), then the mouse received a water droplet. A False Alarm (FA) occurred when the mouse licked the waterspout after a non-target. Discrimination sensitivity was quantified as d’ = z(H)-z(FA). Each behavioral session lasted 30-60 minutes. Analysis of behavioral data was done using custom MATLAB software.

### Widefield Calcium Imaging

#### Chronic Window Implantation

Mice were given an intraperitoneal injection of dexamethasone (5mg/kg) at least 1 hour prior to surgery to prevent inflammation and edema. Mice were deeply anesthetized using isoflurane (5% induction, 0.5-2% for maintenance) and given a subcutaneous injection of cefazolin (500mg/kg). Internal body temperature was maintained at 37.5 C using a feedback-controlled heating blanket. Scalp fur was trimmed using scissors, and any remaining fur was removed using Nair. The scalp was disinfected with alternating swabs of 70% ethanol and betadine. A patch of skin over the temporal bone was removed and the underlying bone cleared of connective tissue using a scalpel. The temporal muscle was detached from the skull, and the skull was cleaned and dried. A thin layer of cyanoacrylate glue (VetBond) was applied to the exposed skull surface and a 3D printed stainless steel head-plate was affixed to the midline of the skull. A circular craniotomy (3 mm diameter) was made over the auditory cortex where the glass imaging window was implanted. The space between the glass, the brain, and the skull was sealed with an optically transparent silicone elastomer (Kwik-Sil). Dental cement (C&B Metabond) was used to cover the entire head-plate and the edges of the glass and the skull were then sealed. Finally, the entire implant except for the imaging window was coated with black dental cement created by mixing methyl methacrylate with iron oxide powder to reduce optical reflections. Meloxicam (0.5mg/kg) was given subcutaneously as a post-operative analgesic.

#### Image Acquisition

Widefield images were acquired as previously described^43,44^. To examine periodotopy and tonotopy, we performed widefield imaging experiments in the auditory cortex of Thy1-GCaMP6s × CBA mice. Awake mice were placed into a 3D-printed plastic tube and head-restraint system. Blue excitation light was shone by an LED (473 nm) through an excitation filter (470 nm) and directed into the cranial window. Emitted fluorescence from neurons was collected through a 4x objective (Thorlabs), passed through a longpass filter (cutoff: 505 nm), followed by a bandpass emission filter (531 nm) attached to a CMOS camera. Images were acquired at a rate of 5 Hz and resolution of 512×512 pixels using ThorCam software (Thorlabs). After acquiring an image of the cortical surface, the focal plane was advanced to approximately 750 μm below the surface for imaging neural activity.

#### Acoustic Stimuli

Acoustic stimuli were presented to mice from an ES1 free field speaker, as described above. To measure tonotopy, the frequency of each pure-tone was randomly selected from 10 equiprobable values (2-45 kHz, 2 tones per octave). To measure periodotopy, the F0 of PTs, HCTs, AMNs, and IRNs were randomly selected from 13 equiprobable values (20-1280 Hz, 2 F0s per octave). Trials began with a 1 s silent period, followed by a 0.5 s stimulus, and ended with a 2 s silent period. Each stimulus was repeated 20 times per experiment, with inter-stimulus intervals randomized at 6, 7, or 8 s. Mice were imaged once per day, and only one type of stimulus was presented per !~20-minute imaging session.

#### Image Registration

Widefield calcium imaging data were preprocessed in MATLAB. For each session, cortical surface images were averaged to provide a high-contrast template for alignment. Fluorescence movies were registered frame-by-frame to the surface reference using rigid translation-based alignment (*imregtform*, Image Processing Toolbox, MATLAB). Registered movies were processed with a 16-pixel spatial filter, downsampled by a factor of four, half-wave rectified, and a variance mask was applied to suppress low-signal regions.

#### Analysis of Periodotopic Gradients

For each stimulus condition, imaging frames were aligned to stimulus onset and averaged across repetitions to obtain a mean time series for every pixel. For each pixel, ΔF/F traces were calculated by finding the average fluorescence during the silent pre-stimulus baseline period (F), subtracting that value from subsequent time points until 1 s after onset, and then dividing all time points by the baseline fluorescence. To identify auditory responses, we retained traces within 90% of the maximum ΔF/F observed at each pixel (ΔF/F_90_). Each pixel was assigned a preferred frequency (for pure-tones) or preferred F0 (for complex tones), defined as the median of the F0 or frequency that evoked ΔF/F_90_ responses. The median was chosen instead of the maximum to provide a stable estimate when several adjacent stimuli produced similar responses. This procedure yielded tonotopic and periodotopic maps.

To segment individual gradients within periodotopic and tonotopic maps^45^, first the ΔF/F_90_ images were smoothed with a 5×5 moving average filter, and local maxima within 10-pixel neighborhoods were identified as candidate gradient centers. The corresponding tonotopic or periodotopic values of these local maxima were then subtracted from all other values in the map. A value of 1 was then added to all pixels to ensure positive values, which facilitates finding reversal points in gradients. Then the tonotopic and periodotopic maps were smoothed with a 2×2 moving average filter to regularize maps. From each center, radial trajectories were traced using Bresenham’s line algorithm^46^, with angles discretized in 0.25° steps. Along each trajectory, reversals in preferred frequency or F0 values were defined as local peaks, and the first reversal from the center was taken as a gradient boundary. Reversal points that were more than 2 standard deviations (STDs) from the mean boundary distance were removed. Boundaries were extended to include pixels within three pixels of a reversal and converted into polygonal masks. Masks were further smoothed with a 5-pixel moving average. Smoothing changes the binary mask into a continuous distribution that tapers to zero beyond the original boundary. To account for this blurring, we removed regions of the smoothed mask that fell below 1 STD from the mean value of the smoothed mask. To reduce redundancy, we took the intersection of the masks with >75% overlap.

Within each segmented gradient, gradient vectors were computed on a per-pixel basis as the change in preferred frequency or F0 (on a log2 scale) divided by cortical distance, oriented along the unit vector between points^45^. Tonotopic and periodotopic vectors were calculated as the normalized mean of gradient vectors. Segemented tonotopic gradients were used to identify auditory cortical fields, based on expected field locations and gradient directions in cortical space^47^. We quantified periodotopic structure using three metrics: vector strength (VS) was defined as the mean resultant length of these vectors, gradient size (Size) was defined as the percentage of auditory cortex occupied by a given gradient, and gradient angle (θ) was computed from the summed vector components.

### Statistical Analyses

#### Non-Parametric Bootstrap t-test

To determine significantly different values between experimental conditions, we used a non-parametric bootstrap t-test, as previously described^43^. Given 2 datasets, A and B, having sample sizes of n and m, respectively, we tested A and B against the null hypothesis that they were drawn from a common distribution. The hypothesis test began by taking the absolute value of the observed difference of means, Δμ, between A and B. Next, we created the null distribution by pooling the individual values of A and B. Two sample sets, A* and B*, of size min(n,m), were randomly selected (with replacement) from the null distribution. The test statistic, Δμ*, was computed from the absolute value of the difference of the means obtained from the A* and B* sample sets. We repeated the random selection of A* and B* from the null distribution and the calculation of Δμ*, 10,000 times, to form a bootstrap distribution of Δμ*. A was taken to have a statistically significant different mean than B, if Δμ* was greater than Δμin less than 5%, 1%, or 0.01% of the 10,000 bootstrapped values. This would mean that the probability was <5%, 1%, or 0.01% that samples in A and B came from a common distribution. All mean values are reported with standard errors of the mean (SEMs). Where noted in the text, multiple comparisons across groups were corrected with a Bonferroni procedure.

#### Circular Statistics

Gradient angles were defined relative to the cortical rostrocaudal and dorsoventral axes. An Omnibus test was used to determine whether angle distributions deviated from uniformity.

#### Linear Regression of Behavioral Pitch Discrimination on Structural Features of Periodotopy

To assess the relationship between behavioral F0 discrimination performance and structural properties of periodotopy, we performed repeated-measures standardized linear regressions. Behavioral sensitivity (d′) and periodotopic predictors (VS, Size, and θ) were measured in separate cohorts of mice. The fixed set of predictor values (one per stimulus condition) served as the within-subject repeated-measures factor. All variables were z-scored prior to analysis, and regressions were fit separately for each behavioral mouse tested in figure 1d. This yielded per-mouse slope coefficients and coefficients of determination (R^2^), which were then averaged across animals. Statistical significance of mean slopes was assessed using the bootstrap procedure described above, with Bonferroni correction for multiple predictors.

### Data Availability Statement

Data will be made available on a public repository and associated with a DOI.

## Notes

### Competing Interest Statement

The authors have declared no competing interest.

